## Supplementary figures and images for "Human airway macrophages are metabolically reprogrammed by IFN-γ resulting in glycolysis dependent functional plasticity"

### Supplemental figure 1

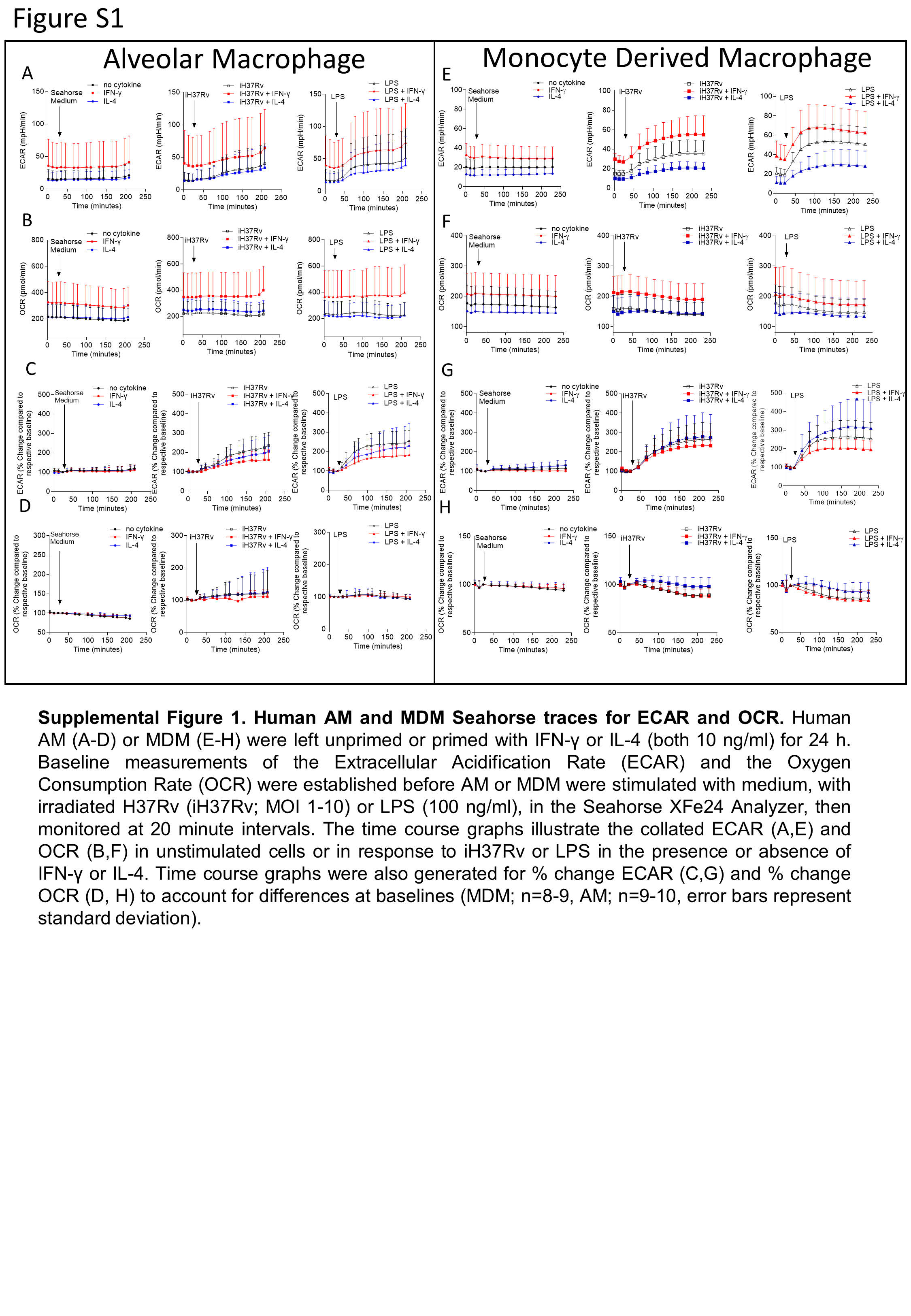

### Supplemental figure 2

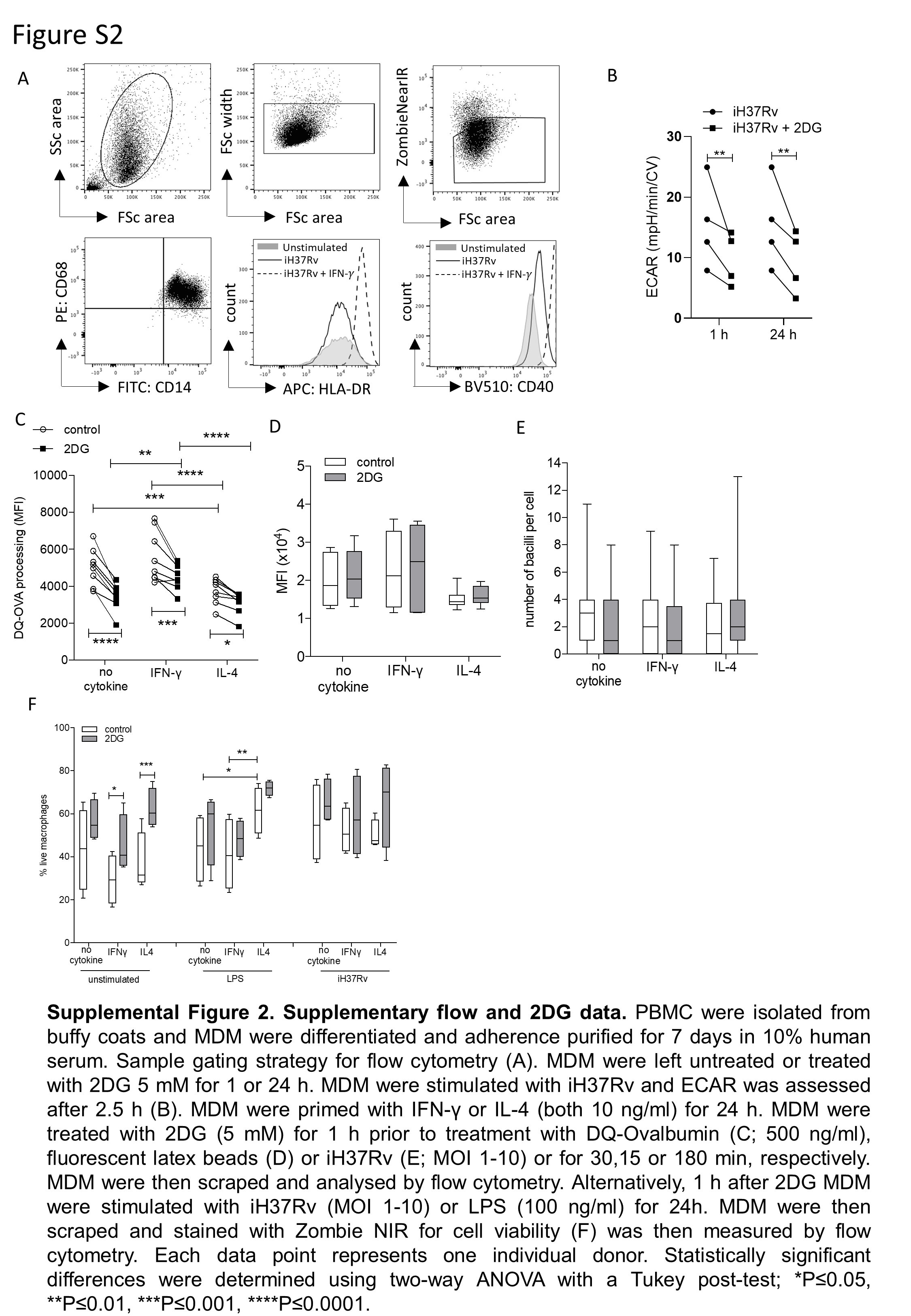
